# Counter-stimuli Inhibit GRPR Neurons via GABAergic Signaling in the Spinal Cord

**DOI:** 10.1101/489831

**Authors:** Rita Bardoni, Devin M. Barry, Hui Li, Kai-Feng Shen, Joseph Jeffry, Qianyi Yang, Antonella Comitato, Yun-Qing Li, Zhou-Feng Chen

## Abstract

A myriad of counter-stimuli, including algogens and cooling, could inhibit itch sensation; however, the underlying molecular and neural mechanisms remain poorly understood. Here, we show that the spinal neurons expressing gastrin releasing peptide receptor (GRPR) primarily comprise excitatory interneurons that receive direct and indirect inputs from C and Aδ fibers and form contacts with projection neurons expressing the neurokinin 1 receptor (NK1R). Optical or chemogenetic activation of GRPR neurons evokes itch behavior that is partly dependent on NK1R activation. Importantly, we show that noxious or cooling counter-stimuli inhibit the activity of GRPR neurons via GABAergic signaling. By contrast, capsaicin, which could evoke a mix of itch and pain sensations, could exert both excitatory and inhibitory effects on GRPR neurons. These data strengthen the role of GRPR neurons as a key circuit for itch transmission and illustrate a spinal mechanism whereby counter-stimuli inhibit itch by suppressing the function of GRPR neurons.

**Highlights:**

1. Activation of GRPR neurons evokes itch and is dependent upon NK1R activation
2. GRPR neurons receive both direct and indirect inputs from C/Aδ fibers
3. Counter-stimuli inhibit GRPR neurons via GABAergic signaling
4. Increased excitability of GRPR neurons in chronic itch condition

## Introduction

Pain and itch information is conveyed by distinct yet interacting neuronal pathways in sensory neurons and spinal cord to the brain (Akiyama and Carstens, 2013; Barry et al., 2017; Bautista et al., 2014; Braz et al., 2014; LaMotte et al., 2013). GRPR is a member of the mammalian homologs of the bombesin-receptor family and plays an important role in a number of physiological functions, including itch sensation (Barry et al., 2017; Jeffry et al., 2011; Jensen et al., 2008). GRPR and GRPR neurons in the superficial dorsal horn of the spinal cord are important for transmitting itch information from the skin to the spinal cord (Akiyama et al., 2014; Barry et al., 2018; Barry et al., 2017; Liu et al., 2011; Sun and Chen, 2007; Sun et al., 2009). GRPR neurons are subject to a myriad of regulations through cross-talk with other G protein coupled receptors, such as MOR1D, an isoform of mu opioid receptor (Liu et al., 2011), or 5HT1A, a serotonin receptor(Zhao et al., 2014a), KOR, a kappa opioid receptor(Munanairi et al., 2018), resulting in activation (MOR1D), facilitation (5HT1A) or attenuation (KOR) of the activity of GRPR neurons.

GRPR neurons may form contacts with NK1R neurons which project to the parabrachial nucleus (PBN) and the spinothalamic track (STT) neurons to relay itch information in mice (Akiyama et al., 2015; Cameron et al., 2015; Mu et al., 2017). Although some information about GRPR neurons have been obtained using indirect approaches or transgenic mice which may not fully recapitulate *Grpr* expression in the spinal cord(Aresh et al., 2017; Wang et al., 2013), direct characterization of molecular, anatomical and electrophysiological properties of GRPR neurons are yet to be achieved. For example, although ablation of GRPR neurons abolished itch but not pain transmission(Sun et al., 2009), whether lack of pain behaviors can be attributable to a compensatory effect is unclear. It is also debatable with respect to the mode of itch transmission, including input and output pathways of GRPR neurons (Barry et al., 2017; Bautista et al., 2014; Braz et al., 2014).

The pain pathway can suppress itch transmission as shown by numerous anecdotal evidence and experimental studies (Akiyama et al., 2011; Ikoma et al., 2006; Ward et al., 1996). Other notable counter-stimuli include cooling (e.g. menthol), which relieves itch via transient receptor potential cation channel subfamily M member 8 (TRPM8) in DRG neurons (Bromm et al., 1995; Palkar et al., 2017). It has been shown that counter-stimuli inhibit itch through GABAergic neurons in the spinal cord (Kardon et al., 2014). However, the precise molecular mechanisms and the identity of the dorsal horn neurons inhibited by GABAergic signaling remain elusive.

We used a combination of anatomical tracing, optogenetics, chemogenetics, behavioral analysis, and electrophysiology to characterize the properties of GRPR neurons. Our studies reveal previously unknown features of GRPR neurons and shed important light onto how itch is inhibited by counter-stimuli.

## Results

### GRPR neurons in the dorsal horn and SpVc are interneurons and form synaptic contacts with projection neurons

To directly test whether GRPR neurons are interneurons or projection neurons, we performed retrograde tracing of projection neurons by injecting the retrograde fluorescent tracer Fluoro-Gold (FG) into the thalamus or PBN of GRPR-eGFP mice followed by double immunohistochemistry (IHC) staining as described (Fig. 1. a-d, h-k) (Zhao et al., 2014b). GRPR neurons are mainly distributed in laminae I and II (Fig. 1, green). Although not all *Grpr* neurons expressing eGFP, our previous studies found that all eGFP neurons analyzed express *Grpr* as validated by single cell RT-PCR and their responses to GRP (Zhao et al., 2014a). These eGFP neurons were primarily located in the superficial dorsal horn (laminae I-II), both medially and laterally(Zhao et al., 2014a). FG-labeled lamina I neurons were found predominantly in the spinal trigeminal nucleus caudalis (SpVc) and upper cervical segments of the spinal cord after FG injection into the thalamus (Fig. 1. e-g, red), while after PBN injection, the majority of FG neurons were found in lumbar segments (Fig. 1. l-n, red). Of 150 sections examined from different segments of the spinal cords and SpVc of mice (n = 15) that were injected with FG into thalamus or PBN, none of the eGFP neurons were co-labeled with FG. Consistent with the fact that the majority of NK1R neurons are PBN-projecting neurons in mice (Cameron et al., 2015), eGFP was not co-labeled with NK1R nor with NK1R/FG double-labeled neurons (Fig. 1. o-r). However, numerous eGFP contacts were observed with NK1R neurons, suggesting that itch information from GRPR neurons is transmitted through NK1R neurons (Fig. 1 s, t).

**Fig. 1.**
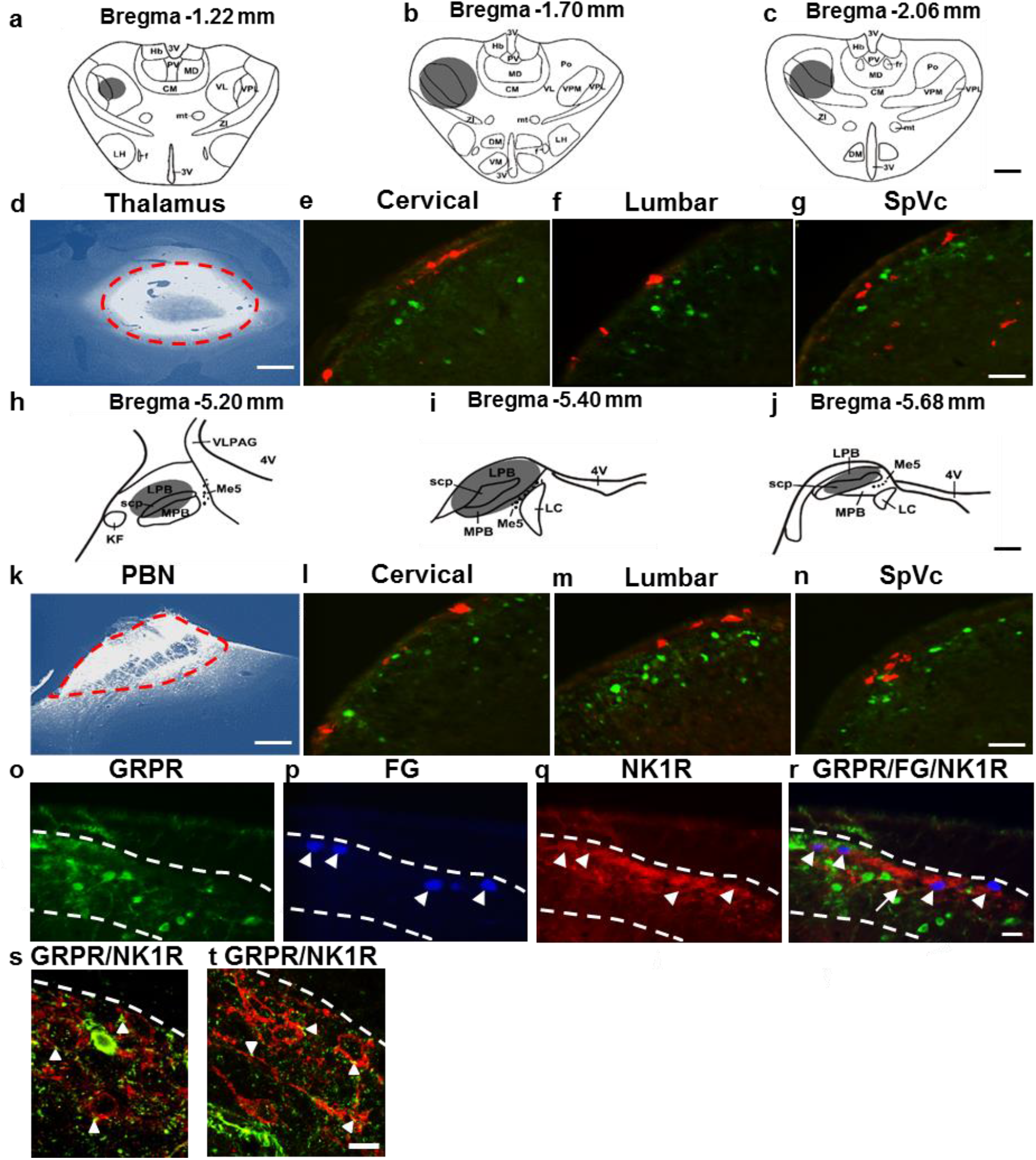
GRPR^+^ neurons in the spinal cord dorsal horn and SpVc are interneurons. **a-c** Diagrams show FG injection sites (grayed areas) in the thalamus. **d** FG (bright white) injection site in the thalamus was circled in red dashed line. **e-g** There was no GRPR (GFP, green) and FG (red) double-labeled cells in the dorsal horn of the cervical spinal cord **(e)**, lumbar spinal cord **(f)** and SpVc **(g)** in GRPR-eGFP mice. **h-j** the grayed areas indicate the injection site **(h)** and diffused regions **(i, j)** of FG after PBN injection. **k** Red dashed line defines the border of injection site of FG in PBN. **l-n** Double staining in the dorsal horns of cervical spinal cord **(l)**, lumbar spinal cord **(m)** and SpVc **(n)** in *Grpr*-eGFP mice showed that *Grpr* (GFP, green) neurons were not FG (red) projection neurons to PBN. **o-r** GRPR neurons **(o)** were not co-labeled with FG (**p**, arrowheads), NK1R (**q**, arrowheads), and FG/NK1R double-labeled neurons (**r**, arrowheads). s, **t** GRPR terminals (green) make contacts (arrowheads) with NK1R neurons (red) in lamina I of spinal dorsal horn. Scale bars, 600 μm in A-D, H-K; 25 μm in E-G, L-R; 10 μm in **s** and **t**.

We next examined whether GRPR neurons form direct connection with PBN projecting neurons using double immuno-electron microscopic (Immuno-EM) for GRPR and FG in the lumbar cord. Terminals of GRPR neurons identified by the silver enhanced nanogold particles formed asymmetric synapses with FG dendritic profiles revealed by the immunoperoxidase reaction product (Supplementary Fig. 1).

### Optogenetic or chemogenetic activation of GRPR neurons evokes itch behavior that is partly dependent on NK1R activation

Our initial finding that GRPR neurons are interneurons that make connections with NK1R neurons prompted us to functionally investigate GRPR-NK1R connectivity in itch transmission. We first generated *Grpr*-improved Cre recombinase knock-in (*Grpr*^iCre^) mice (Supplementary Fig. 2a). We validated the efficacy of iCre expression by double *in situ* hybridization (ISH) in *Grpr*^iCre^ spinal cord dorsal horn and found that nearly all *iCre*^+^ neurons co-expressed *Grpr* (115/128 *iCre*^+^ neurons) and, likewise, almost all *Grpr*^+^ neurons co-expressed *iCre* (115/126 *Grpr*^+^ neurons) (Fig. 2a-c). We also confirmed functional activity of i*Cre* through intraspinal injection and expression of a Cre-dependent adeno-associated virus (AAV) expressing eYFP and observed eYFP^+^ neurons in laminae I and II, consistent with expression patterns found previously for *Grpr* (Supplementary Fig. 2b). Next we characterized *Grpr*^iCre^ expression in the dorsal horn by crossing *Grpr*^iCre^ mice with the tdTomato flox-stop reporter line Ai9 (Madisen et al., 2010) to generate mice with tdTomato expression in *Grpr*^+^ neurons (*Grpr*^tdTom^) and performed IHC with several markers (Fig. 2d). *Grpr*^tdTom^ neurons received many GRP^+^ contacts in laminae I and II (Fig. 2e). Consistent with previous results of GRPR-eGFP expression (Zhao et al., 2014b), most *Grpr*^tdTom^ neurons co-expressed the excitatory marker Lmx1b (84.7 ± 1.4%, 283/334 neurons)(Cheng et al., 2005), but showed less co-expression with the inhibitory marker Pax2 (17.2 ± 2.6%, 54/314 neurons) and almost no co-expression with NK1R (1.9 ± 0.6%, 6/330 neurons)(Fig. 2G-I), which supports tracing experiments that GRPR neurons are interneurons.

**Fig. 2.**
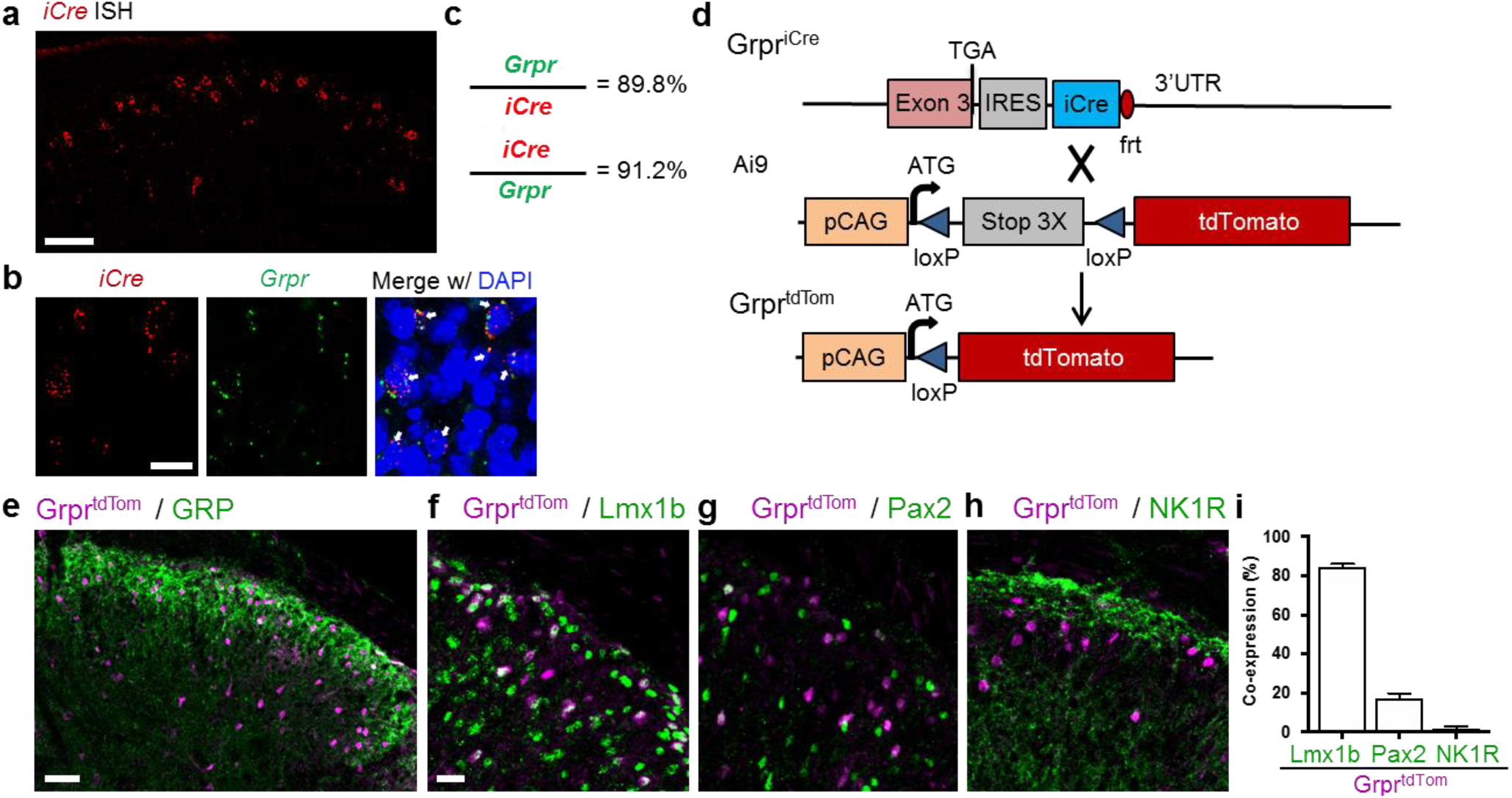
Molecular characterization of spinal *Grpr*^iCre^ neurons. **a** RNAscope *in situ* hybridization (ISH) expression pattern of *iCre* in cervical dorsal horn from Grpr^iCre^ mice. (scale bar, 100 μm). **b** Representative images of ISH of iCre (red) and *Grpr* (green) mRNA expression in superficial dorsal horn of transverse cervical sections. Arrows indicate double-positive neurons. Blue represents DAPI nucleic acid stain (scale bar 20 μm). **c** Percentage of i*Cre*- or *Grpr*-expressing neurons that express *Grpr* or i*Cre*, respectively. n = 3 mice and 15 sections. **d** Schematic of *Grpr*^iCre^ mating with Ai9 reporter line to produce *Grpr*^tdTom^ mice. **e-h** IHC images of *Grpr*^tdTom^ dorsal horn neurons and (**e**) GRP (scale bar, 50 μm), (**f**) Lmx1b (scale bar 20 μm, (**g**) Pax2 or (**h**) NK1R. **i** Percentage of co-expression of Lmx1b, Pax2 or NK1R in *Grpr*^tdTom^ neurons. n = 3 mice and 10 sections per marker. Data are represented as mean ± S.E.M.

To address whether normal pain behavior of mice with spinal ablation of GRPR neurons could be due to compensatory effect(Sun et al., 2009), optogenetics was used to test the role of activation of GRPR neurons in the spinal cord dorsal horn in itch and pain behaviors. To selectively activate GRPR neurons, *Grpr*^iCre^ mice were crossed with the flox-stop channel rhodopsin-eYFP (ChR2-eYFP) line Ai32 (Madisen et al., 2012) to generate mice with ChR2-eYFP expression in *Grpr* neurons (*Grpr*^ChR2^) and implanted fiber optics onto the spinal column at the cervico-thoracic or lumbar regions for blue-light stimulation (Fig. 3a, b). We first tested a brief stimulation time (5 s) at varying frequencies (5, 10, 20 or 30 Hz) to determine the behavioral responsiveness of *Grpr*^ChR2^ or littermate control (*Grpr*^WT^) mice. *Grpr*^ChR2^ mice with cervico-thoracic stimulation showed both scratching and biting responses directed toward the nape, forelimb or upper torso, consistent with the cervico-thoracic (C-T) sensory dermatomes in the periphery (Fig. 3c)(Supplementary Video 1), whereas *Grpr*^WT^ mice showed no response. Induced behavior can be inhibited by i.p. morphine (data not shown)(Akiyama et al., 2010), suggest that induced behavior reflects itch not pain. Consistent with calf model for itch(LaMotte et al., 2011), lumbar (L) stimulation resulted in biting responses directed toward the hind limb, lower back, and urogenital regions, again consistent with the lumbar sensory dermatomes (Supplementary Fig. 3a)(Supplementary Video 2). Behavior response durations increased with stimulation frequency with 30 Hz behavior showing similar duration to the 5 s stimulation time (~4.67 s for C-T; ~5.5 s for L) (Supplementary Fig. 3b, c). The onset of behavior responses became more rapid with decreased latency, increasing frequency and reached a floor effect at 20 Hz for both C-T and L stimulation (~1.2s for C-T; ~0.7s for L) (Supplementary Fig. 3d, e). However, the offset of behavior responses was relatively consistent across frequencies (0.7-1.2s for both C-T and L) (Supplementary Fig. 3d, e). Stimulation response percentages were 75% or more at 10, 20 or 30 Hz for C-T stimulation and 80% or more for L stimulation (Supplementary Fig. 3f, g).

**Fig. 3.**
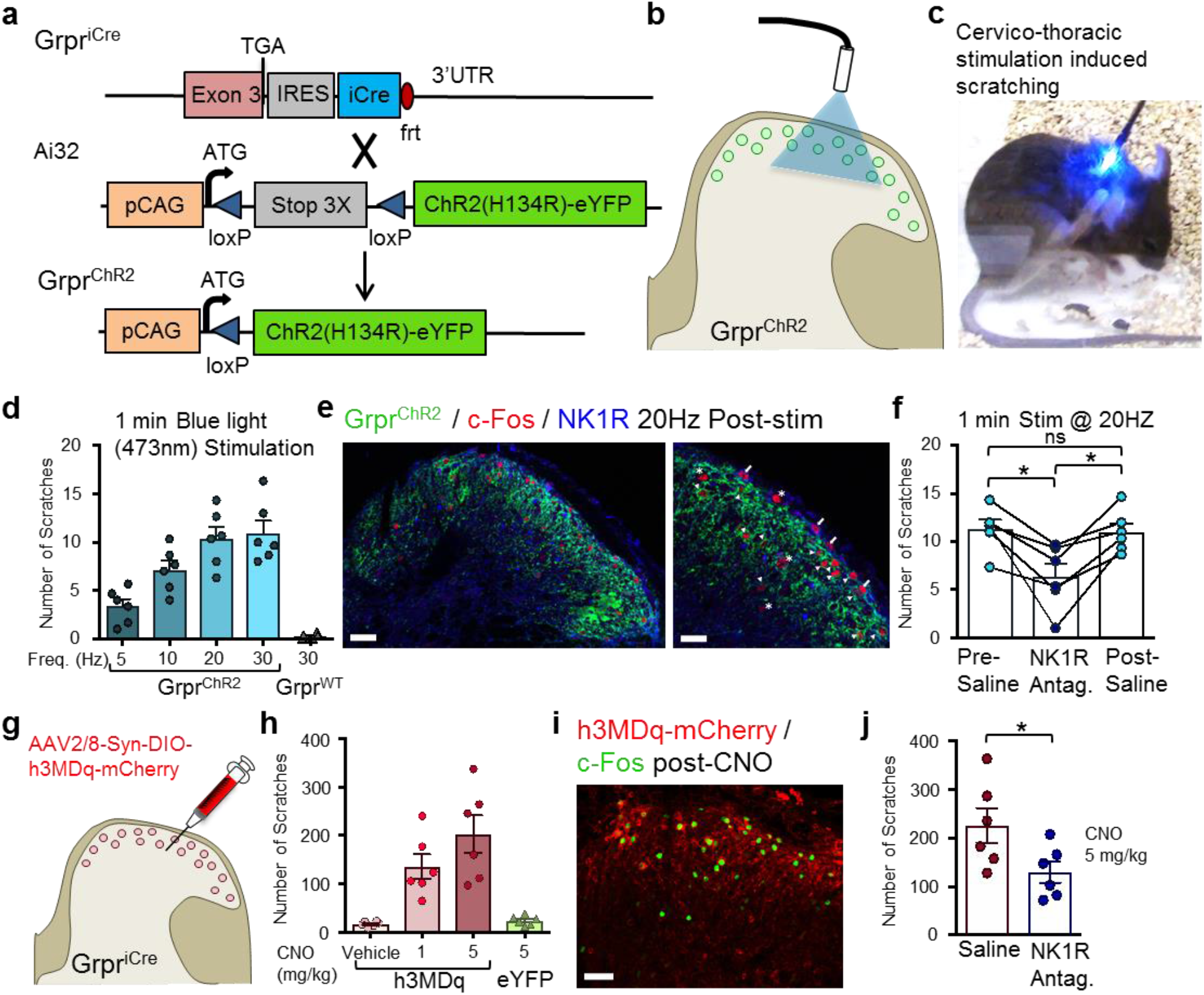
Optogenetic or chemogenetic activation of spinal GRPR neurons induces itch behavior that can be attenuated with spinal NK1R antagonism. **a** Schematic of *Grpr*^iCre^ mating with Ai32 line to produce *Grpr*^ChR2^ mice. **b** Diagram of blue light (473 nm) stimulation of GRPR neurons in spinal cord. **c** Snapshot of Grpr^ChR2^ mouse with scratching behavior induced by stimulation of cervico-thoracic GRPR neurons. **d** Number of scratches induced by 1 min. stimulation at 5, 10, 20 or 30 Hz in *Grpr*^ChR2^ mice or 30 Hz in *Grpr*^WT^ control mice. **e** IHC images of ChR2-eYFP, c-Fos and NK1R following 20 Hz stimulation in spinal cord of *Grpr*^ChR2^ mice. Arrows indicate c-Fos/NK1R double-positive neurons. Arrowheads indicate c-Fos/ChR2-eYFP double-positive neurons. Asterisks indicate c-Fos positive-only neurons. Left image, scale bar 50 μm. Right image, scale bar 20 μm. **f** Number of scratches induced by 1 min. cervico-thoracic stimulation at 20 Hz in *Grpr*^ChR2^ mice with intrathecal (i.t.) injection of saline in pre-test, NK1R antagonist L733,060 (20 μg) or saline in post-test. n = 6 mice, *p < 0.05; one-way RM ANOVA with Tukey multiple comparisons post-hoc analysis. **g** Diagram of intraspinal injection of AAV2/8-Syn-DIO-h3MDq-mCherry in *Grpr*^iCre^ mice. **h** Number of scratches induced by vehicle or CNO (1 or 5 mg / kg, intraperitoneal) in *Grpr*^iCre^ mice with intraspinal injection of h3MDq or control eYFP. **i** IHC image of h3MDq-mCherry and c-Fos following CNO induced scratching behaviors in *Grpr*^iCre^-hM3Dq injected mice. **j** Number of scratches induced by CNO (5 mg/kg) in *Grpr*^iCre^-hM3Dq mice with i.t. injection of saline or L733,060. n = 6 mice, *p < 0.05; Student’s unpaired *t* test. Data are represented as mean ± S.E.M.

To further investigate the effect of GRPR neuron activation in transmitting itch, we used a longer 1 min stimulation period in *Grpr*^ChR2^ C-T implanted mice and found the number of scratches increased with increasing frequency stimulation (Fig. 3d). Following stimulation, c-Fos expression was detected in many *Grpr*^ChR2+^ neurons (Fig. 3e, arrowheads) and NK1R neurons (Fig. 3e, arrows) of the spinal cord dorsal horn, as well as some c-Fos neurons with no apparent labeling (Fig. 3e, asterisks), whereas red-light stimulated control showed no c-Fos expression (Supplementary Fig. 3h). To test whether NK1R signaling is involved in GRPR neuron-mediated itch, an NK1R antagonist (L-733,060, 20 μg), which was previously shown to reduce GRP-induced itch (Akiyama et al., 2014), was pre-injected intrathecally (i.t.) for 20 Hz C-T stimulation and significantly attenuated scratching responses when compared to pre and post saline i.t. injections in *Grpr*^ChR2^ mice (Fig. 3f). Similarly, pre-injection of the NK1R antagonist significantly reduced itch responses for L stimulation of *Grpr*^ChR2^ mice (Fig. S3i).

We next tested the effect of chemogenetic activation of GRPR neurons on itch behavior. A viral vector expressing Cre-dependent G_q_-coupled designer receptors exclusively activated by designer drugs (DREADDs), hM3D_q_ (AAV2/8-Syn-DIO-h3MDq-mCherry), was injected unilaterally into the upper cervical (C2-C5) spinal cord of *Grpr*^iCre^ mice (Fig. 3g). Clozapine-*N*-oxide (CNO, 1 or 5 mg/kg) induced robust scratching behaviors to the nape, head and face, whereas vehicle or AAV-DIO-eYFP control mice injected with CNO induced little scratching (Fig. 3h). Following CNO injection and behavior, c-Fos expression was detected in many *Grpr*-hM3D_q_ neurons as well as some hM3D_q_-negative neurons (Fig. 3i). Lastly, blockade of NK1R signaling by i.t. injection of the NK1R antagonist significantly attenuated scratching behavior induced by CNO in *Grpr*^iCre^-hM3D_q_ expressing mice (Fig. 3j). Taken together, activation of GRPR neurons, using both optogenetic and chemogenetic approaches, induces itch behavior that is dependent, in part, on downstream activation of NK1R in the spinal cord.

### Characterization of GRPR neuron membrane properties

To characterize the properties of GRPR neurons, electrophysiological recordings were obtained from a total of 230 GRPR-eGFP neurons in the spinal cord slices from P16-P25 mice. Action potential firing patterns were determined from a sample of 39 GRPR neurons, by recording, in current clamp, the responses to injections of depolarizing current. Most neurons (56.4%) exhibited a delayed firing pattern, when current steps were applied from a membrane potential of about −80 mV (Fig. 4a, d). This pattern is characterized by a delay in the generation of the first action potential, that is larger than the average interspike interval (Fig. 4b). Other subpopulations of GRPR neurons showed a tonic (23.1%) or a phasic (15.4%) firing pattern (Fig. 4a, d). The tonic pattern is characterized by an action potential discharge that persists during the whole current step and often decreases in frequency. The delay of the first action potential is comparable to the average interspike interval (Fig. 4b). Neurons showing the phasic pattern fire only at the beginning of the current step, with a variable number of action potentials. Only two neurons exhibited a single spike pattern.

**Fig. 4.**
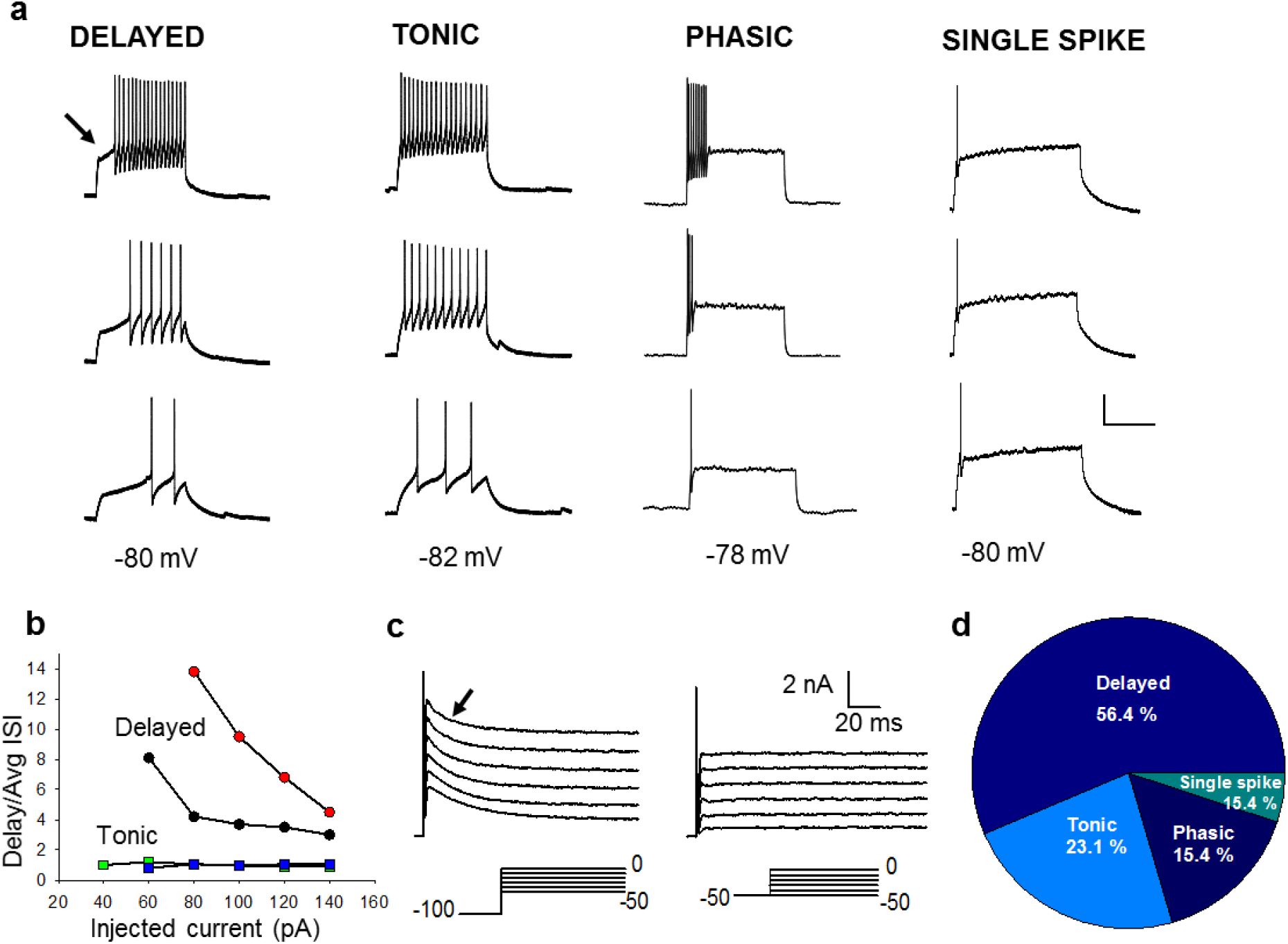
Discharge patterns observed in GRPR neurons. **a** Representative current clamp recordings obtained from GRPR neurons, by holding the membrane potential around −80 mV. The 3 traces for each firing type represent (starting from the lower trace): response to the first current step able to induce action potentials (rheobase) and responses to 2 stimuli above threshold. Scale Bar: 30 mV, 200 ms. **b** Graph representing the criterion used to discriminate between delayed and tonic firing neurons. Delayed firing neurons showed a delay to the 1st spike (indicated by the arrow in A) that was disproportionately long compared with the average interval between spikes (Avg ISI), even at large depolarizing current steps (circles, example of 2 neurons). Tonic firing neurons had delays to the 1st spike comparable to the average ISI (squares, 2 neurons). **c** Example of voltage clamp recordings obtained from a delayed firing GRPR neuron, in the presence of 1 μM TTX. When the neuron was held at −100 mV, voltage steps from −50 to 0 mV evoked a transient, potassium A-type current (marked by the arrow). When the same neuron was held at −50 mV, the A current was absent. **d** Proportions of neurons exhibiting the different firing types, from a total sample of 39 GRPR neurons.

Passive and active membrane properties of delayed, tonic and phasic firing GRPR neurons were determined (Supplementary Table 1). Notably, tonic firing neurons showed lower rheobases and more negative action potential thresholds than delayed firing neurons, consistently with previous studies (Punnakkal et al., 2014).

Voltage clamp recordings performed from delayed firing neurons showed the presence of a transient, voltage-dependent A current, that was activated by holding the cells at negative potentials (−100 mV) and applying depolarizing voltage steps (Fig. 4c). Activation of the A current is responsible for the delayed firing pattern observed in most GRPR neurons at −80 mV. When these delayed firing neurons were maintained in current clamp at −60/−65 mV, membrane potentials at which the A current is inactivated, they showed different firing patterns, with a prevalence of the tonic firing Supplementary Fig. 4). This is consistent with a recent study(Aresh et al., 2017), showing that GRPR neurons predominantly exhibit a tonic firing pattern at the resting potential.

To confirm the expression and functionality of GRPR on GRPR neurons, we tested the response of 22 neurons to GRP. Application of 1 μM of GRP induced a slow inward current in 73% (16/22) of GRPR neurons, held at −50 mV in voltage clamp. A second application of GRP, performed on a subpopulation of 7 responsive neurons, elicited in 3 cells an inward current of smaller amplitude (11.5 ± 6.8 pA versus 22.6±10.1 pA at the first GRP application). The suppression of an inward rectifier K^+^ current and/or the activation of a non-selective cation conductance could contribute to the GRP-generated current (Supplementary Fig. 5) (Hermes et al., 2013).

### GRPR neurons receive direct inputs from primary afferents

Previous studies showed that abundant GRP fibers are present in the dorsal horn (Sun and Chen, 2007; Takanami et al., 2014; Zhao et al., 2013) and Immuno-EM studies confirmed that GRP fibers form contacts with dendrites of GRPR neurons(Barry et al., 2016). To characterize the primary afferent fibers synapsing onto GRPR neurons, we stimulated the dorsal root attached to the slice and recorded evoked excitatory postsynaptic currents (EPSCs) from GRPR neurons (Fig. 5). Stimulus intensities were determined in separate sets of experiments, by stimulating one end of the dorsal root and recording compound action potentials from the other end (Fig. 5a). Intensities of stimulation ranged between 10 and 25 μA for Aβ fibers, 25 and 100 μA for Aδ fibers, and between 200 and 500 μA for C fibers. These values are consistent with those reported by previous studies performed in mice of comparable age (Daniele and MacDermott, 2009).

**Fig. 5.**
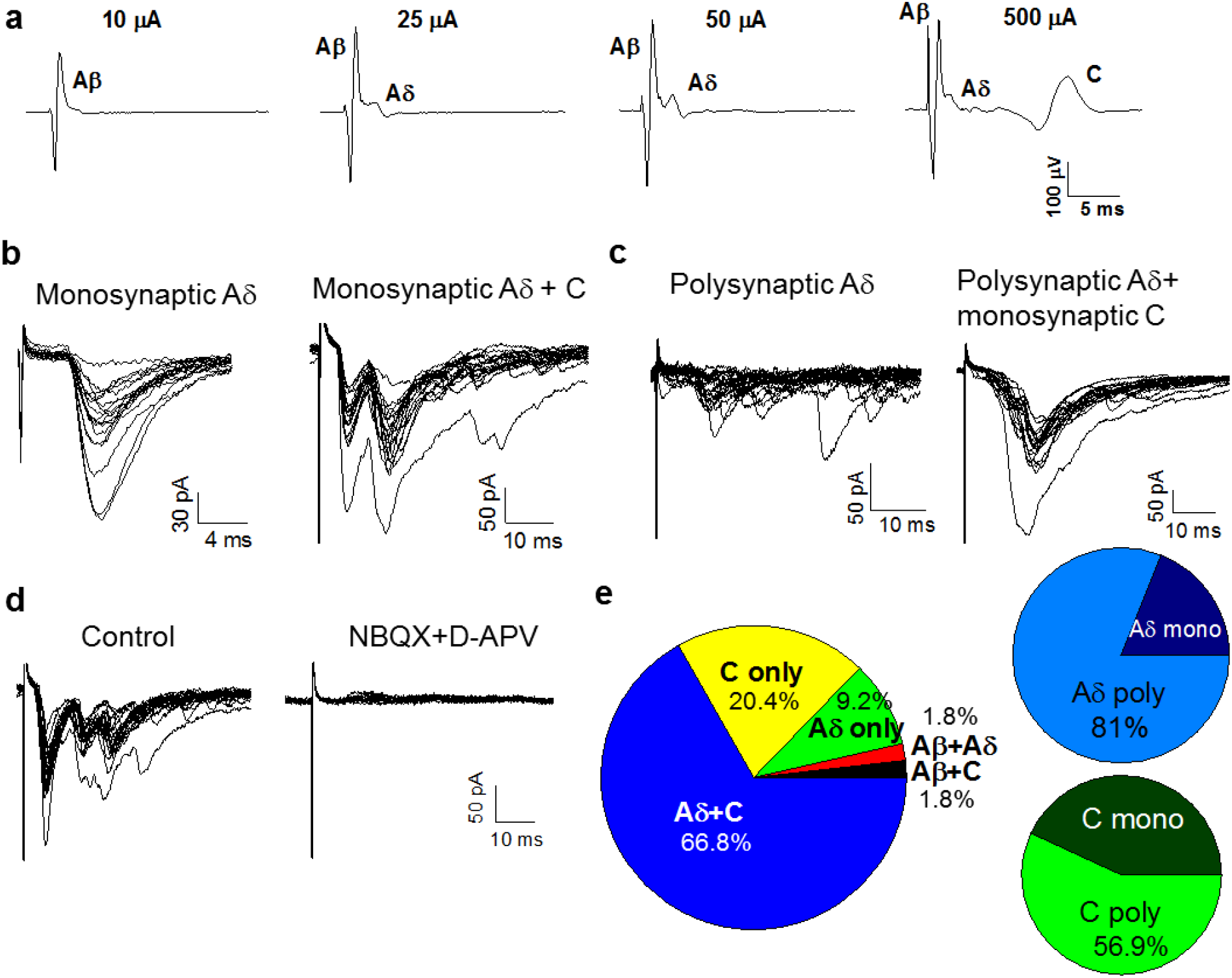
Characterization of excitatory synaptic inputs to GRPR neurons. **a** Compound action potentials (CAPs) recorded from a lumbar dorsal root of a *Grpr*-eGFP mouse. CAPs were evoked by stimulating the dorsal root with a suction electrode at different stimulus intensities. Labelling indicates the voltage peaks corresponding to the different afferent fibers. **b** Example of evoked EPSCs recorded from a GRPR neuron, at different stimulus intensities and frequencies. An Aδ monosynaptic input is evoked at 50 μA, showing no failures at 20 Hz. By increasing the stimulus intensity to 500 μA, a monosynaptic C input becomes apparent. **c** Example of evoked EPSCs, recorded from a different GRPR neuron, using a similar protocol as in B. This neuron received a polysynaptic Aδ input, that failed at 2 or 10 Hz. By stimulating at higher intensities (500 μA) a monosynaptic C input was recruited. **d** Evoked EPSCs were mediated by glutamate ionotropic receptors (AMPA and NMDA), since they were completely blocked by 10 μM NBQX and 50 μM D-APV (n=10). **e** Characterization of the primary afferent inputs mediating evoked EPSCs in GRPR neurons (n=54). In most neurons, EPSCs were mediated by both Aδ and C fibers. Polysynaptic inputs were predominant for both Aδ and C fibers. Only a very small proportion of synapses could be classified as Aδ-mediated (all polysynaptic).

Of 54 GRPR neurons, the majority of cells received synaptic input from both Aδ and C fibers (66.8%), while a small proportion exhibited EPSCs mediated by Aδ or C fibers only (Fig. 5b, c and e). EPSCs mediated by Aδ were mostly polysynaptic (81%), while almost half of the C fiber mediated responses were monosynaptic (43%). Aδ fiber mediated EPSCs (of polysynaptic nature), evoked at intensities lower than 25 μA, were observed in only 2 of the 54 neurons tested, indicating that GRPR neurons mainly receive high threshold primary afferent inputs. Other 4 cells, showing polysynaptic EPSCs evoked at 25 μA, were classified as receiving Aδ inputs, although at this stimulus intensity some additional Aδ fibers could have been also recruited (Fig. 5a). Being polysynaptic connections, it was not possible to apply the high frequency stimulation protocol to distinguish the 2 types of fibers. EPSCs evoked on GRPR neurons by dorsal root stimulation were completely blocked by co-application of the AMPA receptor antagonist NBQX and the NMDA receptor antagonist D-APV, showing that they are mediated by glutamate receptors (Fig. 5d). Since the EPSCs were evoked at −70 mV, they were mainly mediated by AMPA receptors, while the NMDA mediated component was negligible.

Stimulation of Aδ and C fibers was also able to evoke inhibitory postsynaptic currents (IPSCs) on GRPR neurons held at −10 mV (Fig. 6a). Similar to the EPSCs, most neurons exhibited evoked IPSCs mediated by both Aδ and C fibers (15 out of 25 neurons tested, Fig. 6b). Neurons showing IPSCs mediated by Aδ fibers include also 3 cells where IPSCs were evoked at 25 μA, an intensity also compatible with Aδ stimulation (see above). Application of bicuculline produced more than a 90% block of the evoked IPSCs, showing that they were mainly mediated by GABA_A_ receptors (Fig. 6c).

**Fig. 6.**
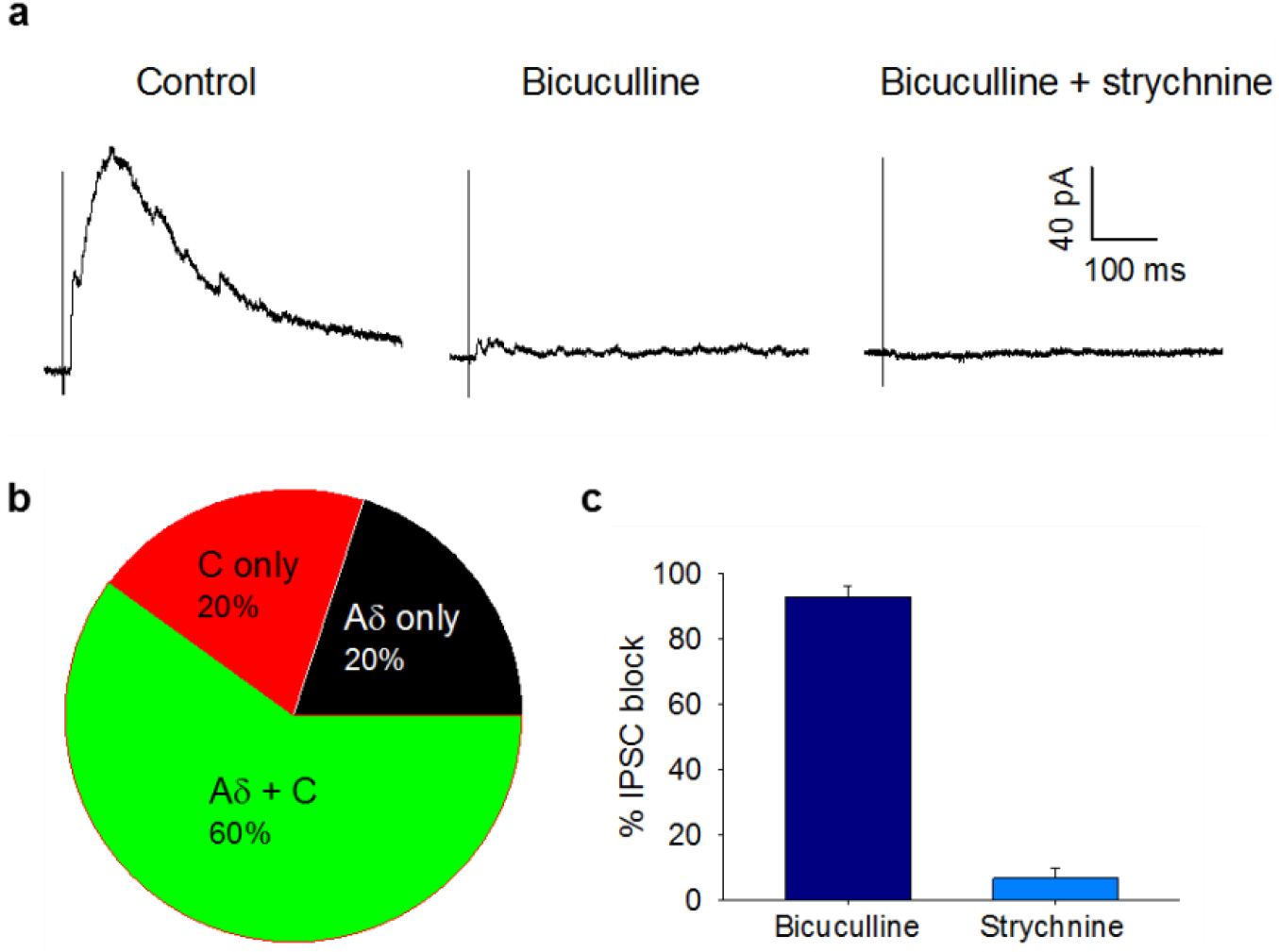
Characterization of inhibitory synaptic inputs to GRPR neurons. **a** Example of evoked IPSCs recorded from a GRPR neuron held at −10 mV. The IPSC in control shows a faster component mediated by Aδ fibers (evoked by stimulating at 50 μA) and a slower current, mediated by C fibers (evoked at 500 μA). 10 μM bicuculline blocked almost completely the IPSC. Application of bicuculline plus 300 nM strychnine caused a complete block. **b** Characterization of the primary afferent inputs mediating evoked IPSCs in GRPR neurons (n=25). In most neurons, IPSCs were mediated by both Aδ and C fibers. **c** Percentage block of the evoked IPSCs, observed in presence of 10 μM bicuculline and 300 nM strychnine (n=10). GABA_A_ receptors mediate most of the IPSCs recorded from GRPR neurons. Data are represented as mean ± S.E.M.

### Inhibition of GRPR neurons by counter-stimuli

The observation that evoked IPSCs on GRPR neurons are mediated by high threshold primary afferents suggests that these neurons are inhibited by nociceptive inputs. To test this hypothesis, we examined the effect of the counter-stimuli capsaicin, allyl isothiocyanate (AITC, a key component of mustard oil) and menthol on spontaneous IPSCs (sIPSCs) recorded from GRPR neurons. Upon application of menthol (500 μM), we observed a significant increase of sIPSC frequency in 26.3% of GRPR neurons, with an average 3.3-fold increase (Fig. 7a, b and i). AITC (500 μM) significantly increased the sIPSC frequency in 21.6 % of neurons, with an average 5.3-fold increase (Fig. 7c, d and i). An increase of sEPSC frequency was rarely observed in both menthol and AITC (Fig. 7j). Remarkably, application of capsaicin (1 μM) affected both sEPSCs and sIPSCs of GRPR neurons (Fig. 7e-j), with a potent effect (23.5-fold increase) on sEPSC frequency in the large majority of cells tested (91.6%). By contrast, sIPSC frequency was increased 4.6-fold in 34.3 % of the recorded neurons. These results confirm that counter-stimuli are effective in activating inhibitory spinal interneurons, producing an increase of the inhibitory tone in GRPR neurons.

**Fig. 7.**
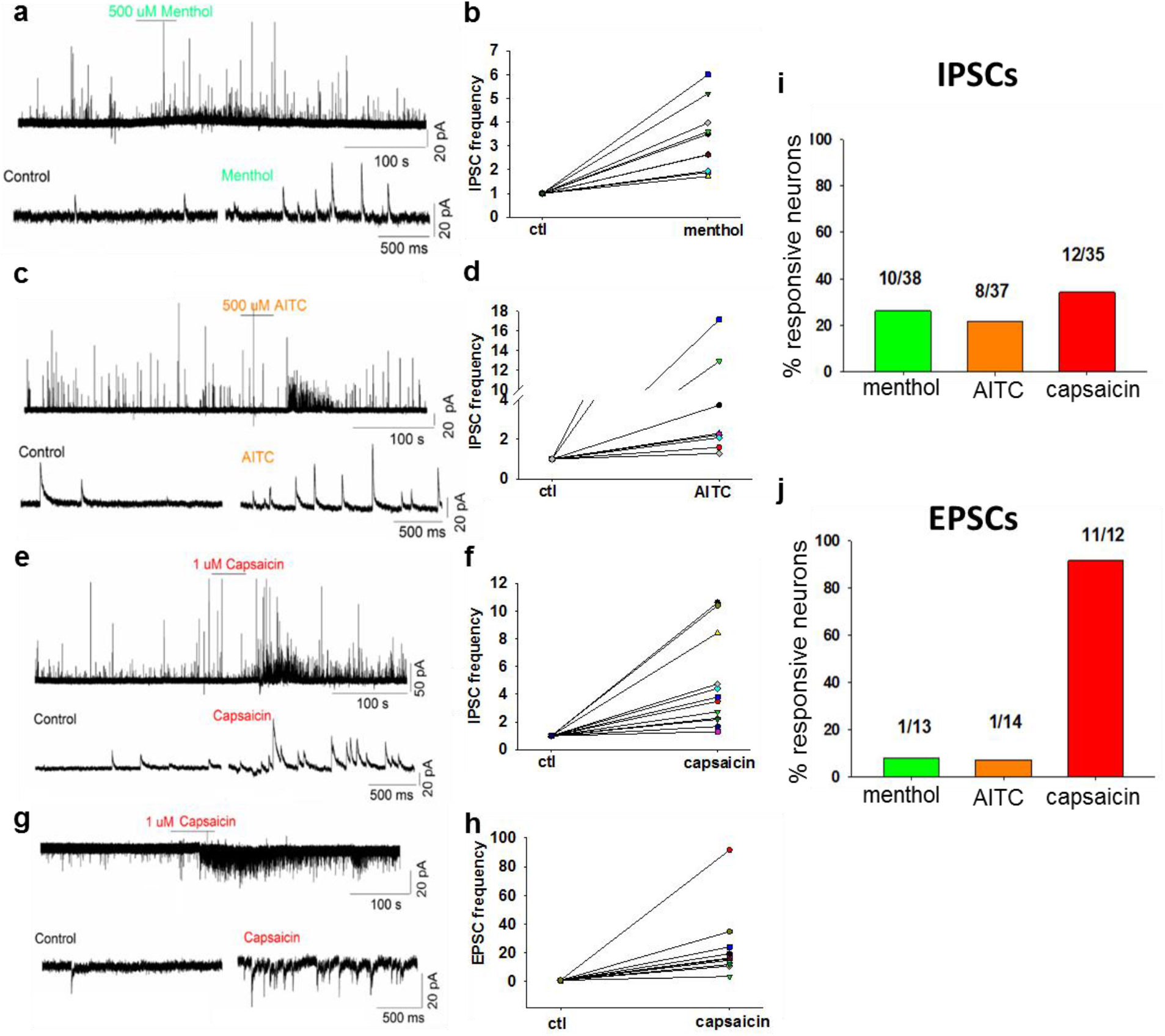
Counter-stimuli increase inhibition onto GRPR neurons. **a, c, e** Example traces of spontaneous IPSCs recording from GRPR neurons, held at −10 mV. One min application of 500 μM menthol (**a**), 500 μM AITC (**c**) or 1 μM Capsaicin (**e**) produced a significant increase of sIPSC frequency in subpopulations of GRPR neurons. Statistical significance (P<0.05) was determined by using the Kolmogorov-Smirnov test on individual neurons. Lower traces depict sIPSCs on an expanded time scale for control and counter-stimuli applications. **b, d, f** Normalized sIPSC frequencies observed in the responsive neurons in the presence of the counter-stimuli. **g** Application of capsaicin induced also a strong frequency increase of sEPSCs, recorded at – 60 mV. **h** Scatter plot of normalized sEPSC frequencies, obtained from a sample of GRPR neurons responsive to capsaicin. **i, j** percentages of GRPR neurons exhibiting an increase of sIPSC or sEPSC frequency upon application of the 3 counter-stimuli. Data are represented as mean ± S.E.M.

### Spontaneous firing of GRPR neurons from BRAF^Nav1.8^ mice

At last, we investigated the excitability of GRPR neurons during chronic itch using BRAF^Nav1.8^ mice, a genetically engineered mouse model in which GRP and GRPR expression is dramatically increased (Zhao et al., 2014a), by performing current clamp recordings. At membrane potentials between −50 and −55 mV, only 1 out of 19 GRPR neurons from w/t mice exhibited spontaneous and repetitive action potential firing. On the contrary, GRPR neurons derived from BRAF^Nav1.8^ mice showed, in the same experimental conditions, a high level of spontaneous synaptic activity, accompanied by spontaneous firing (Supplementary Fig. 6a, b). This increase of excitability was observed in 11 out of 31 neurons tested and was particularly evident during the first 3-5 min of recording, after which a spontaneous hyperpolarization occurred. Accordingly, the average frequency of action potentials fired by GRPR neurons from BRAF^Nav1.8^ mice, during the first minute of recording, was significantly higher than that observed in GRPR neurons from w/t mice, recorded in the same conditions (Supplementary Fig. 6c). These data indicate that chronic itch increases the excitability of GRPR neuron, which could be related to modifications in the synaptic inputs, such as increased GRP release as suggested by increased GRP expression in primary afferents (Zhao et al., 2013).

## Discussion

In this study, using traditional neuroanatomical tracing and immuno-EM approach we demonstrate that GRPR neurons are interneurons that make contacts with NK1R PBN- and STT-projecting neurons. Electrophysiological studies show that the majority of GRPR neurons exhibit, at hyperpolarized potentials, a delayed firing pattern, typical of characteristics of dorsal horn excitatory interneurons previously described (Heinke et al., 2004; Yasaka et al., 2010). Importantly, optogenetic stimulation of GRPR neurons resulted in downstream activation of NK1R neurons and itch-related scratching behavior that was partially dependent upon NK1R activation for itch transmission (Akiyama et al., 2015; Carstens et al., 2010; Mu et al., 2017). Combined with the observation of direct contacts between GRPR neurons and NK1R neurons, these data demonstrate that GRPR neurons function as the last relay station prior to sending the information to NK1R projection neurons.

One important finding is that optogenetic or chemogenetic activation of GRPR neurons in the spinal cord dorsal horn evokes itch-related behaviors, arguing against the possibility that normal pain behavior of mice with ablation of spinal GRPR neurons could be due to compensatory effect(Sun et al., 2009). The finding resembles and complements the phenotype of activation of GRP neurons in sensory neurons(Barry et al., 2018). By carefully differentiating between itch- and pain-related scratching behaviors caused by GRPR neuronal activation, these results significantly strengthen the notion that GRPR neurons are dedicated to itch transmission.

That GRPR neurons receive both direct C/Aδ and indirect high threshold inputs from primary afferents is in line with earlier studies in primates and humans showing that primary afferent pruriceptors are high threshold C/A fibers(Handwerker et al., 1987; Johanek et al., 2008; Ringkamp et al., 2011; Schmelz et al., 1997). It is surprising that we failed to find significant A inputs onto GRPR neurons using transverse spinal cord slices, a preparation that has been shown to be suitable for studying mono- and polysynaptic responses mediated by Aδ fibers (Betelli et al., 2015; Torsney and MacDermott, 2006). However, we cannot exclude the possibility that synaptic circuits activating superficial GRPR neurons (sometimes selected because of the stronger fluorescent signal) were not entirely preserved, thereby contributing to an underestimation of the amount of Aδ inputs received by GRPR neurons in the present study. The fact that EPSCs evoked by the dorsal root stimulation were blocked by NBQX and D-APV demonstrates that fast glutamatergic transmission constitutes an integral mechanism for relaying itch information from primary afferents to GRPR neurons. While a high concentration of GRP can directly evoke spikes on GRPR neurons(Aresh et al., 2017), at a lower dose that is of more physiological relevance, GRP could only depolarize cells, suggesting that GRP *in vivo* may modulate glutamatergic transmission under normal physiological condition (Zhao et al., 2014a).

By directly recording the responsiveness of GRPR neurons to chemical counter-stimuli, we show that mustard oil, capsaicin and menthol inhibited the activity of many GRPR neurons. Capsaicin, mustard oil and menthol, acting on different subsets of primary afferent fibers expressing TPRV1, TRPA1 and TRPM8 receptors, respectively(Bandell et al., 2004; Bautista et al., 2006; Caterina et al., 1997; Jordt et al., 2004), enhance the release of glutamate in the dorsal horn to convey pain or cooling sensations. The observation that over 90% of IPSCs evoked by stimulation of Aδ and C fibers were blocked by bicuculline indicates that fast GABAergic transmission is necessary and sufficient to mediate the overwhelming inhibitory effects of these counter-stimuli on GRPR neurons. This is consistent with a recent study reporting that excitatory interneurons located in laminae I and II are predominantly inhibited by GABA (Takazawa et al., 2017).

Unlike mustard oil, a pure noxious stimulus, topical application of capsaicin can induce a mix of pain and itch sensations (Green, 1990; Green and Shaffer, 1993). Interestingly, we found that capsaicin exhibited dual effect on GRPR neurons: while inhibiting some, it also activates GRPR neurons. These results suggest that capsaicin induces pain by activating non-MrgprA3 TRPV1 nociceptors(Han et al., 2013), which in turns inhibits GRPR neurons, while concurrently inducing itch by activating TRPV1 in pruriceptors(Imamachi et al., 2009) and/or MrgprA3 neurons, which mediates itch via innervating GRPR neurons (Han et al., 2013). Coupled with previous studies, it seems that GRPR neurons receive at least three different kinds of inputs from primary afferents: a direct input from C/Aδ pruriceptors, an indirect glutamatergic input from NMBR neurons (Zhao et al., 2014b), and an indirect inhibitory input from GABAergic neurons, in part mediated by vesicular glutamate transporter (Vglut2) in primary afferents (Lagerstrom et al., 2010; Liu et al., 2010). Therefore, there exit several modes of action of GRPR neurons who activation or inhibition could underlie the responses of itch behavior mediated by various counter-stimuli. Moreover, the observation of an enhanced excitability of GRPR neurons in the spinal cord of BRAF^Nav1.8^ mice is consistent with an up-regulation of GRPR expression in chronic itch conditions (Lagerstrom et al., 2010; Shiratori-Hayashi et al., 2015; Zhao et al., 2013). Under these itch conditions, it is conceivable that some nociceptors in sensory neurons may be sensitized to relay enhanced itch information so that capsaicin induces only itch but not pain sensation.

## Experimental procedures

### Animals

Behavioral tests were carried out on C57BL/6J, BRAF^Nav1.8^ (Zhao et al., 2013), *Grpr*^iCre^, GRPR-eGFP (Stock no. 036178-UCD, MMRRC), Ai9 (Stock no. 007909, Jax mice), and Ai32 (Stock no. 024109, Jax mice) unless indicated otherwise. All mice were housed under a 12 h light/dark cycle. Mice were housed in clear plastic cages with no more than 5 mice per cage in a controlled environment at a constant temperature of ~23°C and humidity of 50 ± 10% with food and water available *ad libitum*. All experiments conform to guidelines set by the National Institutes of Health and the International Association for the Study of Pain and were reviewed and approved by the Animal Studies Committee at Washington University School of Medicine. The Italian Ministry of Health approved all electrophysiology experiments in accordance with the Guide for the Care and Use of Laboratory Animals and the EU and Italian regulations on animal welfare.

### Retrograde tracing

GRPR-eGFP male mice were anesthetized with an intraperitoneal injection of ketamine/xylazine (90 mg/kg; 10 mg/kg) cocktail, injected with buprenorphine (BupSR, 0.5 mg/kg) for analgesia, and fixed in a stereotaxic frame (Stoelting, Wood Dale, IL, USA). An incision was made along the midline of the skull, and a small hole was drilled through the bone over the approximate location of the injection sites. A pulled borosilicate glass pipette with a tip in diameter of 20 μm was back filled with mineral oil and attached to a Nanoject II auto-nanoliter injector. The injector was attached to a manipulator and moved to the coordinates. A solution of 4% FG (Biotium, Hayward, CA, USA) was filled into the pipette. FG (0.15-0.25 μl) was injected into each injection site. For thalamus (ventral posterolateral thalamic nucleus-VPL, ventral posteromedial thalamic nucleus-VPM, posterior thalamic nuclear group-Po, posterior thalamic nuclear group, triangular part-PoT), 0.15 μl FG was injected into site a (AP −1.22, ML ± 1.45, DV −3.30), 0.25 μl into site b (AP −1.70, ML ± 1.60, DV −3.40), and 0.15 μl into site c (AP −2.06, ML ± 1.40, DV −3.30). For PB (lateral parabrachial nucleus-LPBN, medial parabrachial nucleus-MPBN, superior cerebellar peduncle-scp, Kölliker-Fuse nucleus-KF), 0.25 μl of FG was injected into one site (AP −5.20, ML ± 1.25, DV −2.40). The stereotaxic co-ordinates were measured from bregma (AP) and the brain surface (DV). After the incision was sutured, triple antibiotic ointment and lidocaine were applied to the skin. Antibiotics (enrofloxacin, 2.5 mg/kg) with saline were injected subcutaneously to prevent infection and the mouse recovered on a warm pad and was returned to the home cage upon walking. Recovered animals were not neurologically impaired. Mice were anesthetized 5~7 days later with an overdose of ketamine/xylazine cocktail and perfused with 0.1 M PBS and then 4% paraformaldehyde. The brain and spinal cord were removed, post-fixed in 4% paraformaldehyde for 6 h at 4 °C, cryoprotected overnight in 30% sucrose in PBS. Brains and spinal cords were sectioned transversely on a cryostat at 50 μm and 20 μm, respectively, for injection sites observation or immunofluorescent staining.

### Generation of *Grpr*^iCre^ mice

An ~8kb genomic DNA fragment containing *Grpr* exon 3 was PCR amplified from a BAC clone (RP-23, id. 402H23, ThermoFisher Scientific) with a high-fidelity DNA polymerase (CloneAmp HiFi PCR mix, Clontech) and subcloned into pBluescript KS. An IRES-iCre and frt-flanked PGK-Neomycin cassette for positive selection were integrated into the 3’UTR immediately downstream of the Stop codon (TGA) using an In-Fusion® HD Cloning Kit (Clontech). A diphtheria toxin α (DTA) cassette was inserted downstream of the 3’ homology arm of the construct as a negative selection marker. The construct was linearized with SacII and electroporated into AB1 embryonic stem cells. Following negative selection with G418 (200 ng / μL), positive clones were identified by 5’ and 3’ PCR screening. Positive clones were injected into C57Bl/6J blastocysts to generate chimeric mice. After breeding, germ-line transmission was confirmed by genotype PCR, mice were bred to a FRT-deleter line (Stock. No. 007844, Jackson Laboratory) to remove the Neomycin cassette and were subsequently backcrossed into the C57Bl/6J background.

### Spinal fiber optic implantation

Implantation of fiber optics onto the spinal column was performed as previously described with some modification (Christensen et al., 2016). Mice were anesthetized with an intraperitoneal injection of ketamine/xylazine (90 mg/kg; 10 mg/kg) cocktail and injected with buprenorphine (BupSR, 0.5 mg/kg) for analgesia. The cervical or lumbar skin was shaved, disinfected and an incision of the skin was made along the midline of the spine. The spinal column was fixed in a stereotaxic frame using spinal adaptors (Stoelting, Cat. No. 51690). After removal of tissue around and between the vertebrae to expose the spinal cord, a small burr hole was drilled ~0.5-0.8 mm lateral from the midline to one side of the vertebra (C7, C8 or T1 for cervico-thoracic, L3 or L4 for lumbar) with a ~0.5mm drill bit with care taken not to drill into the spinal cord. After cleaning and drying the surface of the vertebra, a custom-made ferrule with ~0.25mm fiber optic tip (200 μm in core diameter, Doric Lenses) was placed at the burr hole using a stereotactic holder and super glue gel with accelerant followed by dental cement (Lang Dental) was used to secure the fiber-optic ferrule onto the vertebra. The skin was closed with nylon sutures and triple antibiotic ointment and lidocaine were applied to the skin. Antibiotics (enrofloxacin, 2.5 mg/kg) with saline were injected subcutaneously to prevent infection. Animals were allowed to recover on a warming pad and placed in the home cage for 2 weeks before stimulation.

### Optogenetic Stimulation Behavior

7 – 12-week-old *Grpr*^ChR2^ mice and wild-type littermates (*Grpr*^WT^) were used for optical stimulation experiments. One day prior to the experiments, each mouse was placed in a plastic arena (30 X 30 X 30 cm) for 30 min to acclimate. For stimulation, the fiber optic ferrule spinal implant was connected via a ferrule sleeve to a fiber optic cable with commutator (Doric Lenses) that was attached to a fiber-coupled 473 nm blue laser (BL473T8-150FC, Shanghai Laser and Optics Co.) with an ADR-800A adjustable power supply. The animal was allowed to acclimate being tethered to the cable for 30 min prior to stimulation. Laser power output from the fiber optic was measured using a photometer (Thor Labs) and set to 10 mW from the fiber tip (fiber implants were tested and % efficiencies of power was recorded prior to implantation to ensure 10mW final power from tips). An Arduino UNO Rev 3 circuit board (Arduino) was programmed and attached to the laser via a BNC input to control the frequency and timing of the stimulation. For 5 s stimulation, 10 trials (5 s on, 15 s off) were performed for each frequency (5, 10, 20 or 30 Hz) with 5 min break between each frequency. The mean value of the 10 trials for behavior responses for each frequency was used in the results and analysis for each animal. For 1 min stimulation, 3 trials (1 min on, 10min off) were performed at 20 Hz for each animal and intrathecal injections (saline 10 μL or NK1R antagonist L-733,060, 20 μg / 10μL, Tocris Cat. No. 1145) were performed 10 min before stimulation. Pre-saline, NK1R antagonist and post-saline injections and stimulations were performed on 3 consecutive days, respectively. Again, the mean value of the 3 trials for behavior responses was used in the results and analysis. Mice were recorded with a video camera from a side angle and played back on computer for assessments of the number of scratches, as well as time spent biting, by observers blinded to the animal groups.

### Intraspinal Virus injection

Intraspinal injections of AAVs were performed as previously described (Munanairi et al., 2018). *Grpr*^iCre^ mice were anesthetized with ketamine (90 mg/kg) and xylazine (10 mg/kg) intraperitoneally and injected with buprenorphine (BupSR, 0.5 mg/kg) for analgesia. Cervical vertebrae were exposed at C2-C6 and the vertebral column was mounted onto a stereotaxic frame with spinal adaptors. After removal of tissue around and between the vertebrae to expose the spinal cord, the dura was incised with a sharp needle to expose the spinal cord surface. AAV2/8-Syn-DIO-hM3Dq-mCherry (2.0 X 10^13^ vg/mL) or AAV5-Ef1a-DIO-eYFP (5.6 X 10^12^ vg/mL) was injected into the left side of the spinal cord at 3 sites between successive vertebrae at C3-C4-C5 with a Hamilton Neuros-syringe with beveled needle (catalog number: 65458-02, 34 gauge, 20 degree angle). The syringe needle was inserted into the dorsal spinal cord at an angle of ~35 degrees at a depth of ~250 μm to target the superficial dorsal horn. The AAV was injected (~300 nL AAV per injection) at a rate of 50 nL/min with a Stoelting Quintessential Injector (QSI, catalog number: 53311) and the needle was slowly removed 5 min after the injection was complete. The skin was closed with nylon sutures and triple antibiotic ointment and lidocaine were applied to the skin. Antibiotics (enrofloxacin, 2.5 mg/kg) with saline were injected subcutaneously to prevent infection. Animals were allowed to recover on a warming pad and placed in the home cage for 2 weeks to allow for virus expression before CNO injection.

### *In situ* hybridization

ISH was performed as previously described (Munanairi et al., 2018; Wang et al., 2012). Briefly, mice were anesthetized with a ketamine/xylazine cocktail (ketamine, 90 mg/kg and xylazine, 10 mg/kg) and perfused intracardially with 0.01 M PBS, pH 7.4, and 4 % paraformaldehyde (PFA). The spinal cord was dissected, post-fixed in 4 % PFA for 16 h, and cryoprotected in 20% sucrose overnight at 4 °C. Tissues were subsequently cut into 18 μm-thick sections, adhered to Superfrost Plus slides (Fisher Scientific), and frozen at −20°C. Samples were processed according to the manufacturer’s instructions in the RNAscope Fluorescent Multiplex Assay v2 manual for fixed frozen tissue (Advanced Cell Diagnostics), and coverslipped with Fluoromount-G antifade reagent (Southern Biotech) with DAPI (Molecular Probes). The following probes, purchased from Advanced Cell Diagnostics, were used: *Grpr* (nucleotide target region 463-1596; *GenBank*: NM_008177.2) and *iCre* (nucleotide target region 2-1028; *GenBank*: AY056050.1). Sections were subsequently imaged on a Nikon C2+ confocal microscope (Nikon Instruments, Inc.) in three channels with a 20X objective lens. Positive signals were identified as three punctate dots present in the nucleus and/or cytoplasm. For *Grpr/iCre* mRNA co-localization, dots associated with single DAPI stained nuclei were assessed as being co-localized. Images were taken across the entirety of the population of *Grpr* neurons in each spinal cord section. Cell counting was done by a person who was blinded to the experimental design.

### Immunohistochemistry

IHC was performed as previously described (Barry et al., 2016). 1 - 2 h following optogenetic stimulation or chemogenetic activation with CNO, mice were anesthetized (ketamine, 90 mg / kg and Xylazine, 10 mg / kg) and perfused intracardially with PBS pH 7.4 followed by 4% paraformaldehyde (PFA) in PBS. Tissues were dissected, post-fixed for 2-4 h, and cryoprotected in 20% sucrose in PBS overnight at 4°C. Free-floating frozen sections were blocked in a 0.01 M PBS solution containing 2% donkey serum and 0.3% Triton X-100 followed by incubation with primary antibodies overnight at 4°C, washed three times with PBS, secondary antibodies for 2 h at room temperature and washed again three times. Sections were mounted on slides and ~100 μL Fluoromount-G was placed on the slide with a coverslip. The following primary antibodies were used: chicken anti-GFP (1:500, Aves Labs, GFP-1020), rabbit anti-c-Fos (1:1000, Santa Cruz, sc-52, lot #H1414), guinea-pig anti-NK1R (1:500, AB15810, EMD Millipore), rabbit anti-FG (1:5000, AB153, Millipore). The following secondary antibodies were used: Alexa-Fluor 488 conjugated donkey anti-chicken (1:1000, Jackson ImmunoResearch, 703-545-155), Cy3-conjugated donkey anti-rabbit (1:1000, Jackson ImmunoResearch, 711-165-152) and Cy5-conjugated donkey anti-guinea pig 1(:1000, Jackson ImmunoResearch, 703-175-148). Fluorescent Images were taken using a Nikon C2+ confocal microscope system (Nikon Instruments, Inc.).

### Electrophysiology

#### Spinal cord slice preparation

*Grpr*-eGFP mice (P16-P25) were anesthetized with isoflurane and decapitated, the spinal cord and vertebrae were rapidly removed and placed in ice-cold dissecting Krebs’ solution (composition in mM:125 NaCl, 2.5 KCl, 1.25 NaH_2_PO_4_, 26 NaHCO_3_, 25 glucose, 6 MgCl_2_, 1.5 CaCl_2_, and 1 kynurenic acid, pH 7.4, 320 mOsm), bubbled with 95% O_2_, 5% CO_2_. The lumbar spinal cord was isolated, embedded in an agarose block (low melting point agarose 3%, ThermoFisher Scientific, Waltham, USA), and transverse slices (500 μm thick) were obtained using a vibrating microtome (WPI, Sarasota, USA). Slices were incubated in oxygenated incubation Krebs’ solution (same as dissecting but without kynurenic acid) at 35°C for 30 min and then used for recording.

#### Patch-clamp recording and dorsal root stimulation

Patch-clamp recording in whole-cell configuration was performed on visually identified fluorescent *Grpr*-eGFP neurons at room temperature. Neurons were visualized using an Axioskop microscope (Zeiss, Oberkochen, Germany), fitted with Nomarski optics and connected to a CCD camera (Dage-MTI, Michigan City, USA). Slices were perfused at 2 ml/min with recording Krebs’ solution (in mM:125 NaCl, 2.5 KCl, 1.25 NaH_2_PO_4_, 26 NaHCO_3_, 25 glucose, 1 MgCl_2_, and 2 CaCl_2_, pH 7.4, 320 mOsm). Recordings were performed by using thick-walled borosilicate pipettes (3-5 MOhm resistance), filled with a solution having the following composition (in mM): 120 potassium methane-sulfonate, 10 NaCl, 10 EGTA, 1 CaCl_2_, 10 HEPES, 5 ATP-Mg, pH adjusted to 7.2 with KOH, osmolarity 300 mOsm. Recordings in voltage clamp that required holding the neuron at −10 mV were performed by using an intracellular solution having the following composition (in mM): 130 cesium methane-sulfonate, 10 sodium methanesulfonate, 10 EGTA, 1 CaCl_2_, 10 HEPES, 5 lidocaine N-ethyl bromide quaternary salt-Cl, 2 ATP-Mg, pH adjusted to 7.2 with CsOH, osmolarity 300 mOsm. In voltage clamp experiments, junction potential was corrected after recording. Data were recorded and acquired using a Multiclamp 700A amplifier and the pClamp 10 software (Molecular Devices, Sunnyvale, USA). Sampling rate was 10 kHz, and data were filtered at 2-10 kHz.

To evoke postsynaptic excitatory or inhibitory currents (EPSCs or IPSCs), the dorsal root attached to each slice was stimulated using a suction electrode. Stimulus duration was 0.1 ms, stimulus intensities were determined by performing extracellular recordings of compound action potentials from the dorsal root (see Results and Fig. 5A). Monosynaptic vs polysynaptic EPSCs were identified by stimulating the root at high frequency (20 Hz for Aδ, 10 or 2 Hz for Aδ and 1 Hz for C fibers). EPSCs showing no failures during 20 consecutive stimulations were considered monosynaptic.

Neuron resting membrane potential was determined within the first 5 min of recording and neurons that had a resting membrane potential more positive than −50 mV were discarded. Active and passive membrane properties were determined by applying current steps (10 or 20 pA of amplitude) in current clamp, starting from a membrane potential of −60/−65 mV. Membrane resistance was calculated from responses to hyperpolarizing pulses in the linear region of the current-voltage relationship. Rheobase was defined as the current step magnitude required for the minimum number of action potentials.

Drugs were bath-applied for 1 min. All drugs were obtained from Sigma-Aldrich (Saint Louis, USA), except for GRP, that was purchased from Genscript (Piscataway, USA), and tetrodotoxin (TTX) from Tocris (Bristol, UK). Data were analyzed off-line using pClamp10 or MiniAnalysis (Synaptosoft, Decatur, USA). Graphs were obtained using Sigmaplot 11 (Systat software, San Jose, USA).

### Statistical Analysis

Statistical tests are indicated in figure legends when performed. Values are reported as the mean ± standard error of the mean (SEM). Statistical analyses were performed using Prism 7 (v7.0c, GraphPad, San Diego, CA) or Sigmaplot 11. Normality and equal variance tests were performed for all statistical analyses. *P* < 0.05 was considered statistically significant.

## Supporting information

## SUPPLEMENTARY INFORMATION

Supplemental information includes, seven figures, one table and two videos.

## ACKNOWLEDGEMENTS

We thank the Chen laboratory for comments. D.M.B. has been supported by W.M. Keck Fellowship and NIH-NIDA T32 Training Grant (5T32DA007261-23), The project was supported by AR056318-06 (Z.F.C), NS094344 (Z.F.C), DA037261-01A1 (Z.F.C), The National Natural Science Foundation of China (Grants 1771323 and 81620108008) (H.L. and Y.Q.L.), and local grants to R.B. We also thank J.P.Y., and J.Y. for technical support.

## AUTHOR CONTRIBUTIONS

Z.F.C. conceived the project and designed the experiments; R.B., K.F.S., and J.J., performed electrophysiological recording and data analysis. D.M.B. generated *Grpr*^iCre^ line; D.M.B. and K. F. S. performed and analyzed optogenetic and behavioral experiments. H.L. performed and analyzed anatomical tracing and EM experiments, Q.Y. performed and analyzed ISH experiments; R.B., D.M.B., H.L., and J.J. helped with experimental design and data analysis; R. B. and Z.F.C. supervised the project; A. C. contributed to the work. D.M.B., R.B., H.L., and Z.F.C. wrote the manuscript. ^†^ R.B., D.M.B., H.L., K.F.S., and J.J., contributed equally to this work.

---

**Supplementary Figure 1.**
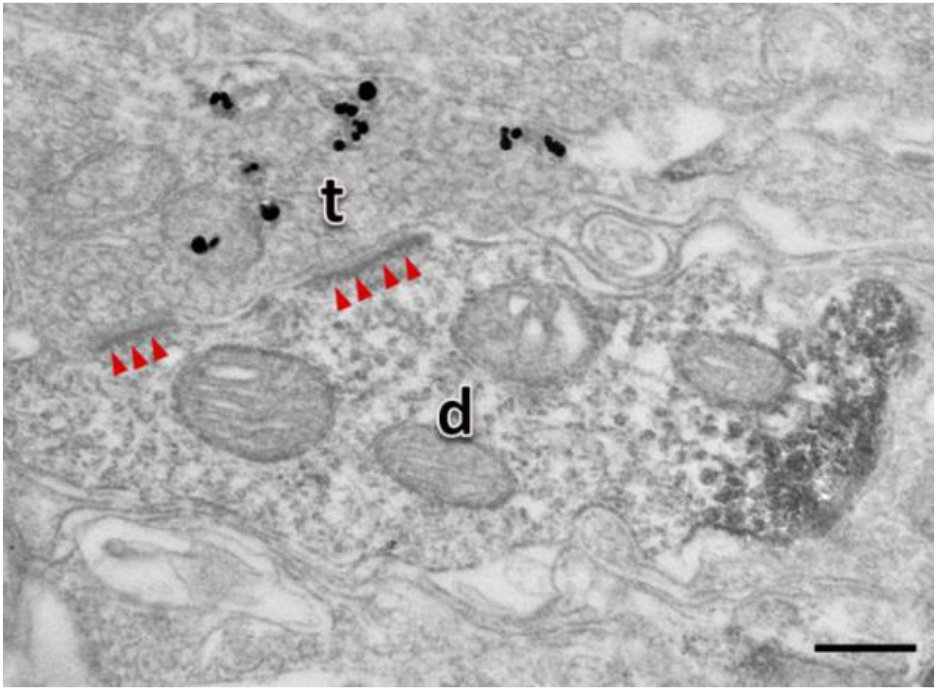
GRPR fibers contacts on PBN-projecting FG retrograde labelled neurons in the superficial spinal dorsal horn. GRPR axon terminal (t; silver grains) makes asymmetric synaptic contact with FG dendritic profile (**d**; DAB reaction products). Arrow heads indicate post-synaptic membranes. Scale Bar, 0.25 μm.

**Supplementary Figure 2.**
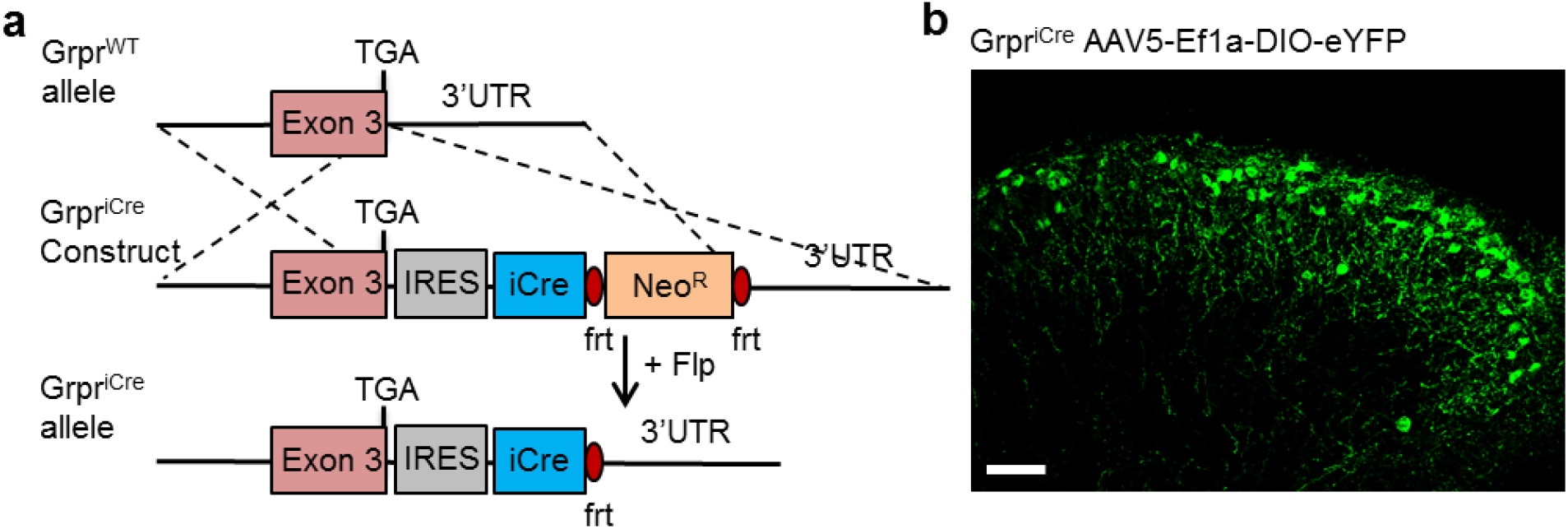
**a** Schematic of targeting strategy for inserting IRES-iCre-Neo cassette into *Grpr* 3’UTR to generate *Grpr*^iCre^ mice. **b** Image of eYFP expression in GRPR dorsal horn neurons of *Grpr*^iCre^ mice injected with AAV-DIO-eYFP. Scale bar, 50 μm.

**Supplementary Figure 3.**
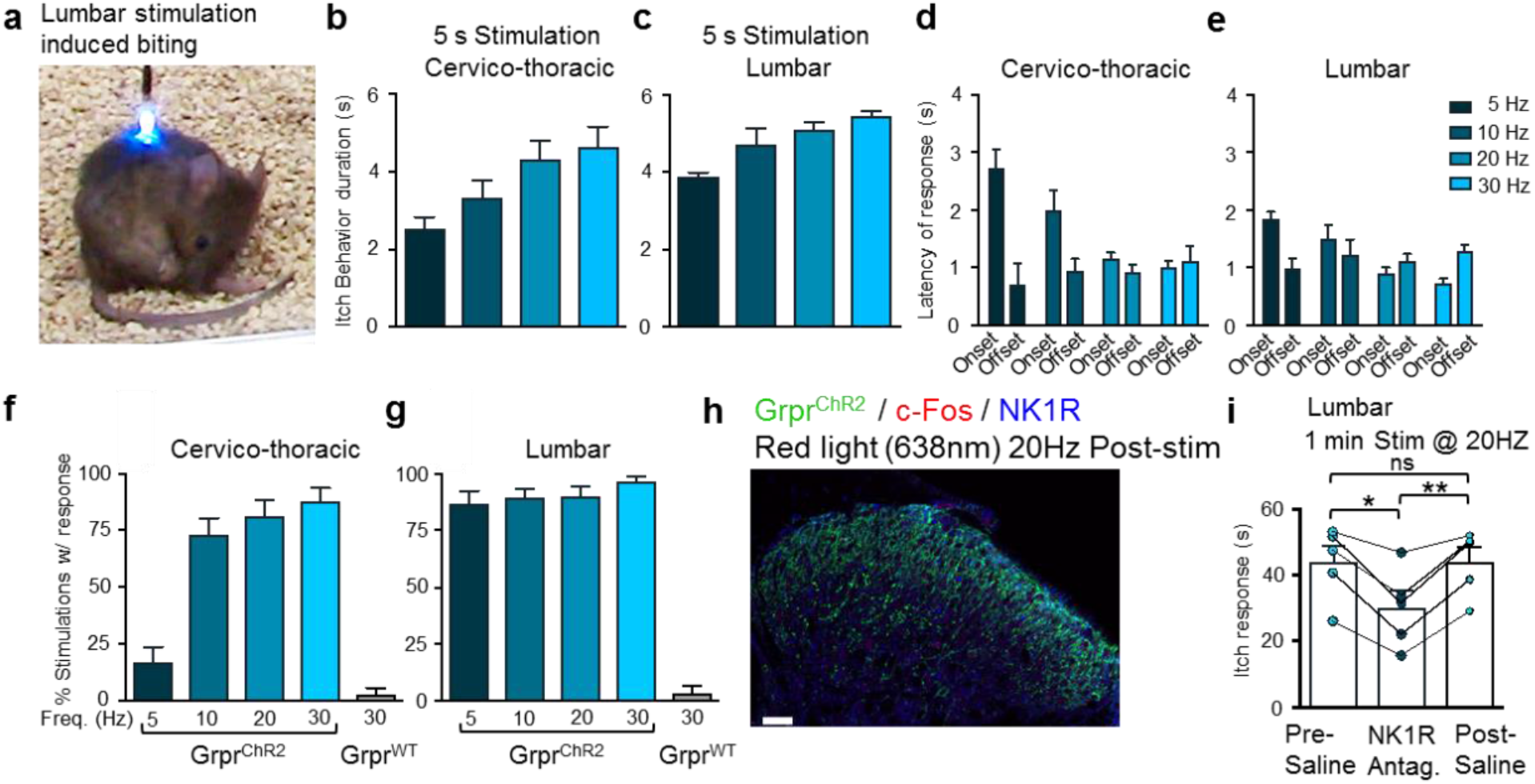
**a** Snapshot of *Grpr*^ChR2^ mouse with biting behavior induced by stimulation of lumbar GRPR neurons. **b, c** Duration of itch behaviors induced by 5 s stimulation of (**b**) cervico-thoracic or (**c**) lumbar neurons at 5, 10, 20 or 30 Hz in *Grpr*^ChR2^ mice. **d, e** Latency of behavior response onset and offset with 5 s stimulation of (**d**) cervico-thoracic or (**e**) lumbar dorsal horn at 5, 10, 20 or 30 Hz in *Grpr*^ChR2^ mice. **f, g** Percentage of stimulations evoking responses at 5, 10, 20 and 30 Hz in in *Grpr*^ChR2^ mice or at 30 Hz in *Grpr*^WT^ mice. **h** IHC image of ChR2-eYFP, c-Fos and NK1R following red light (638 nm) 20Hz control stimulation in spinal cord of *Grpr*^ChR2^ mice. Scale bar 50 μm. **i** Time spent itching induced by 1 min. lumbar stimulation at 20Hz in *Grpr*^ChR2^ mice with intrathecal (i.t.) injection of saline in pre-test, NK1R antagonist L733,060 (20 μg) or saline in post-test. n = 5 mice, * p < 0.05, **p < 0.01; one-way RM ANOVA with Tukey multiple comparisons post-hoc analysis. Data are represented as mean ± S.E.M.

**Supplementary Figure 4.**
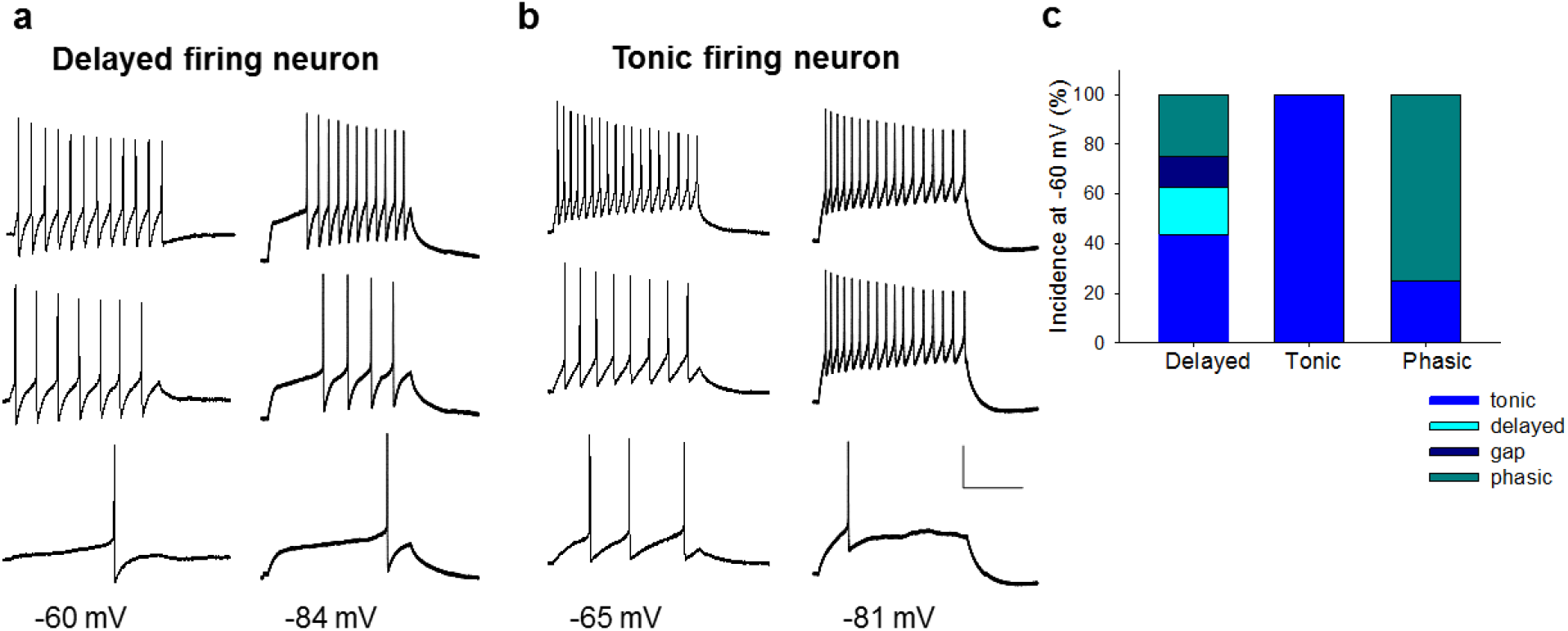
Voltage-dependence of firing patterns in GRPR neurons. **a** Representative current clamp recordings obtained from a neuron exhibiting a delayed firing pattern at −84 mV (right). Holding the neuron near −60 mV, a tonic pattern becomes apparent (left). The 3 traces represent (starting from the lower trace) the response to rheobase and the responses to 2 stimuli above threshold. **b** Sample current clamp recordings obtained from a neuron exhibiting a tonic firing pattern at −81 mV (right). A similar firing pattern is present also at −60 mV (left). Scale Bar: 30 mV, 300 ms. **c** Incidence of the different firing patterns at −60 mV for neurons exhibiting delayed (n=16), tonic (n=8) or phasic patterns (n=4) at −80 mV.

**Supplementary Figure 5.**
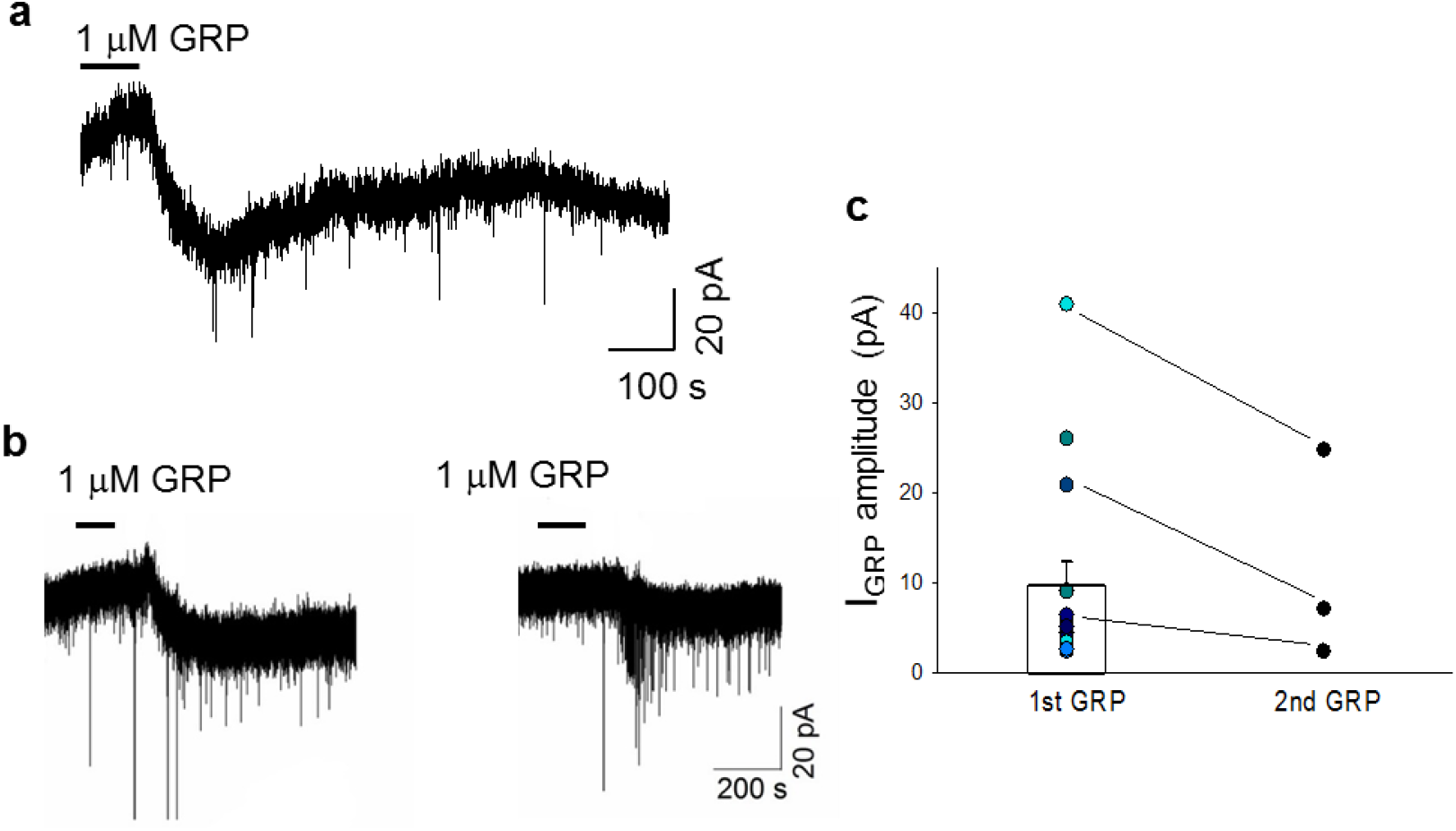
GRP application generates a slow inward current in GRPR neurons. **a** Example trace representing the slow inward current generated by application of 1 μM GRP onto a GRPR neuron held at −50 mV. **b** Currents elicited by 2 consecutive GRP applications onto a different GRPR neuron. **c** Amplitudes of the GRP-induced current recorded from a sample of 16 responsive GRPR neurons. Seven of these neurons were tested for a second application of GRP, that produced an inward current in 3 cells. Data are represented as mean ± S.E.M.

**Supplementary Figure 6.**
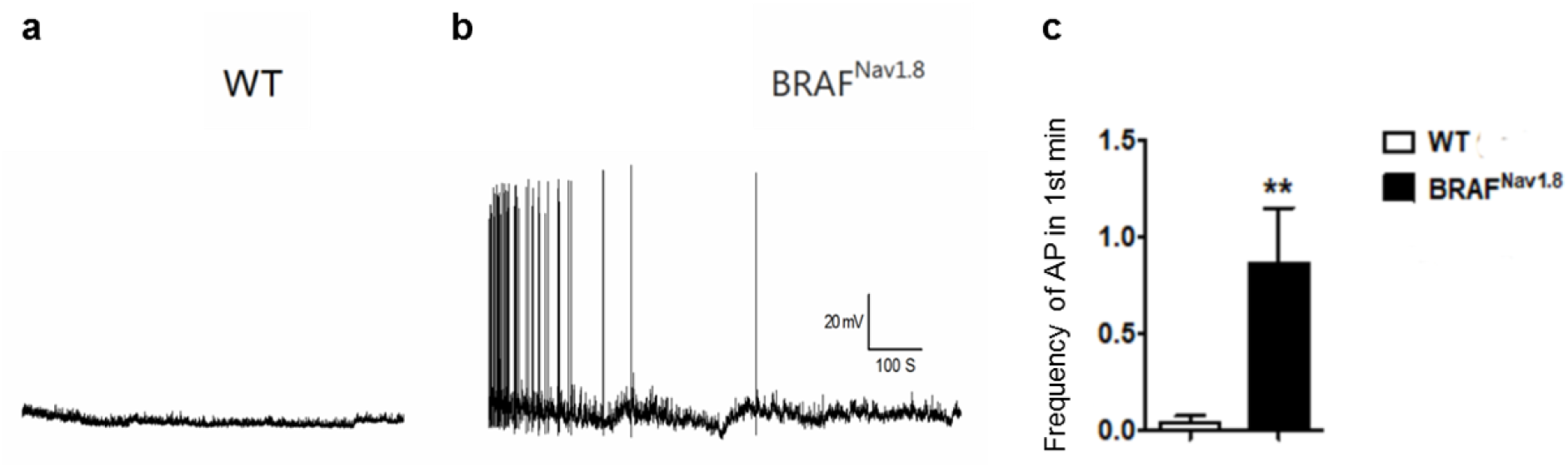
Increased excitability of GRPR neurons in BRAF^Nav1.8^ mice. **a, b** Current clamp recordings at −50/−55 mV obtained from GRPR neurons derived from w/t or BRAF^Nav1.8^ mice. Only GRPR neurons from BRAF^Nav1.8^ mice presented spontaneous, repetitive firing during the early phase of recording. **c** Histogram representing the average frequency of spontaneous action potentials recorded, during the first minute, from the 2 samples of GRPR neurons. Action potential frequency in GRPR neurons from BRAF^Nav1.8^ mice is significantly higher than in w/t mice (t-test, p=0.0085, n=10). Data are represented as mean ± S.E.M.

**Supplementary Table 1.** Passive and active membrane properties of GRPR-eGFP neurons classified according to their firing patterns

|  | **Delayed firing neurons (D)**(n = 13) | **Tonic firing neurons (T)** (n= 7) | **Phasic firing neurons (P)** (n= 5) |
| --- | --- | --- | --- |
| **Resting membrane potential (mV)** | 59.5 ± 1.9 | 56.7 ± 2.5 | 57.8 ± 2.4 |
| **Membrane resistance (MΩ)** | 1020 ± 83.2 | 997 ± 60.1 | 838 ± 83.4 |
| **Rheobase (pA)** | 35.4 ± 4.7<br>* T | 14.3 ± 1.7<br>*D, P | 37 ± 8.6<br>*T |
| **Action potential threshold (mV)** | -35.9 ± 1.1 | -42 ± 1.1 | -37.4 ± 3.6 |

Statistical significance was assessed by one-way ANOVA followed by a Tukey posthoc test. \*, p < 0.05 in the posthoc test, followed by the abbreviation of the group the comparison was made with. n, number of analyzed neurons.

**Supplementary Video 1 Caption.** Cervico-thoracic stimulation (20Hz) induces itch behaviors (scratching in first trial, biting in second trial) in Grpr^ChR2^ mice.

**Supplementary Video 2 Caption.** Lumbar stimulation (20Hz) induces itch behaviors (biting) in Grpr^ChR2^ mice.

